# Variation and evolution of the glutamine-rich repeat region of Drosophila Argonaute-2

**DOI:** 10.1101/048595

**Authors:** William H Palmer, Darren J Obbard

**Affiliations:** Institute of Evolutionary Biology and Centre for infection, Evolution and Immunity, University of Edinburgh, Kings Buildings, West Mains Road, Edinburgh, UK

## Abstract

RNA interference pathways mediate multiple biological processes through Argonaute-family proteins, which bind small RNAs as guides to silence complementary target nucleic acids. In insects and crustaceans, *Argonaute-2* silences viral nucleic acids, and therefore acts as a primary effector of innate antiviral immunity. Although the function of the major of *Argonaute-2* domains, which are conserved across most Argonaute-family proteins, are known, many invertebrate *Argonaute-2* homologs contain a glutamine-rich repeat (GRR) region of unknown function at the N-terminus. Here we combine long-read amplicon sequencing of Drosophila Genetic Reference Panel (DGRP) lines with publicly available sequence data from many insect species to show that this region evolves extremely rapidly and is hypervariable within species. We identify distinct GRR haplotype groups in *D. melanogaster*, and suggest that one of these haplotype groups has recently risen to high frequency in North American populations. Finally, we use published data from genome-wide association studies of viral resistance in *D. melanogaster* to test whether GRR haplotypes are associated with survival after virus challenge. We find a marginally significant association with survival after challenge with Drosophila C Virus in the DGRP, but we were unable to replicate this finding using lines from the Drosophila Synthetic Population Resource panel.

## Introduction

Argonaute proteins are the effectors of eukaryotic RNA interference (RNAi) pathways, using short nucleic acid guide sequences to target complementary sequences for transcriptional or post-transcriptional repression. RNAi-related pathways mediate a diverse range of biological processes, from regulation of developmental genes through miRNAs and endogenous siRNAs, to defence against genomic parasites such as transposable elements via piRNAs (reviewed in Carmell et al, 2002; Meister, 2013). RNAi is also a key line of antiviral defence in plants (Lindbo et al, 1993; Ratcliff et al, 1997), fungi (Segers et al, 2006), ecdysozoan animals such as arthropods and nematodes (Wilkins et al, 2005; Wang et al, 2006), and possibly even in some vertebrate tissues (Li et al, 2013; Maillard et al, 2013). In insects, antiviral RNAi is mediated by an RNA Induced Silencing Complex that contains Agonaute-2 (Ago2). This complex is guided by 21nt siRNAs ‘diced’ from viral replicative intermediates and other dsRNA substrates by Dicer-2 (Okamura et al, 2004; Lee et al, 2004; Wang et al, 2006) and bound to Ago2. Ago2 then uses these siRNAs to target the ‘slicing’ of viral single-stranded RNA, rendering the targeted viral genome or transcript non-functional.

Despite the diverse biological roles played by Argonaute proteins, their structural organisation is generally conserved over deep evolutionary time (Swarts et al, 2014). For example, eukaryotic Argonaute proteins have a PIWI domain that binds and/or ‘slices’ target nucleic acids (Song et al, 2004; Parker et al; 2004), MID and PAZ domains that bind the 3’ and 5’ ends of the small RNA, respectively (Lingel et al, 2003; Ma et al, 2004; Ma et al, 2005; Boland et al, 2010), and an N-terminal domain which is involved in duplex unwinding (Kwak and Tomari, 2012). Nevertheless, in contrast to these highly conserved domains, the N-terminal region of Argonaute proteins tends to be disordered and lack sequence complexity, and is highly variable between species (Hain et al, 2010). This variation is particularly striking in the arthropod antiviral gene, Ago2, where the N-terminal region is often composed of numerous glutamine-rich repeat motifs (‘GRR’; Hain et al, 2010). For example, even between closely related species such as *Drosophila melanogaster* and *D. simulans*, the N-terminal sequence divergence is extensive. In *D. melanogaster*, Ago2 includes one of the most repetitive amino acid sequences in the genome (Jorda and Kajava, 2009), while in *D. simulans* it is markedly different, with only one large duplication of almost the entire N-terminus (Figure 1).

**Figure 1:**
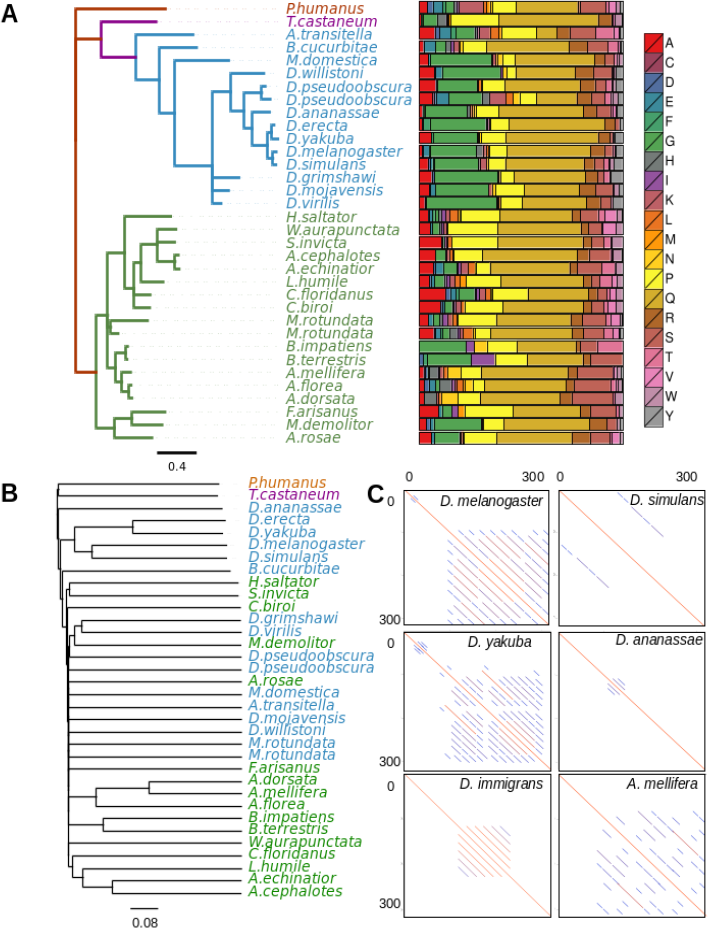
GRR evolves rapidly but maintains similar sequence composition. (A) The gene tree of conserved *Ago2* sequence C-terminal to the GRR, for selected insect species, along with the corresponding amino acid residue composition of the entire GRR for that species. Hymenopteran species are coloured green and dipteran species are coloured blue. Across the insects analysed there is conservation of the residues from which the GRR is composed. (B) Neighbour joining tree drawn from FFP clusters derived from the protein sequence of the entire GRR region, the lack of internal resolution reflects the rapid divergence of the GRR among species. (C) The GRR structure can change rapidly among closely related species. Shown are dotplots for the N-terminal 300 amino acids of Ago2 (plotted against itself) in *D. melanogaster, D. simulans*, *D. yakuba*, *D. ananassae*, *D. immigrans*, and *A. mellifera*. In these dot plots the diagonal line from corner to corner represents the sequence identity to itself, and the successively shorter parallel lines reflect the multiple scales of self-similarity within the sequence.

In *D. melanogaster*, the GRR region is composed of two distinct repeat regions (GRR1 and GRR2; Hain et al, 2010). The most N-terminal, GRR1, is a 6 amino acid imperfect repeat (QQLQQP) present in two to four copies, while GRR2 is a 23 residue imperfect repeat (Figure 2) reported to occur between seven and eleven times in succession in laboratory strains (Hain et al 2010). Although many genetic studies have elucidated the function of Ago2 in *D. melanogaster*, the role of the GRR is still unknown. In other proteins long poly-glutamine tracts have been implicated in increased protein adhesion and protein complex formation, and underlie numerous human diseases (reviewed in Fan et al, 2014). However these are generally long contiguous tracts of glutamine residues, in contrast to the short complex repeat units observed in the Ago2 GRR. Further, *Ago2* GRR deletions appear to have no effect on RISC assembly in *Drosophila* (Liu et al, 2009), suggesting that this domain is not required for binding siRNAs or catalysing target cleavage. The absence of known function makes it difficult to predict which evolutionary forces underlie the observed rapid evolution of the GRR.

**Figure 2:**
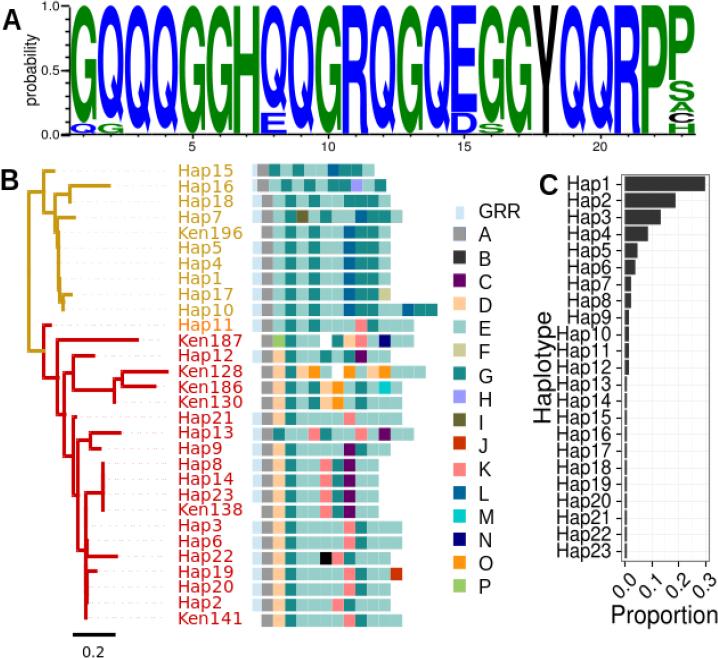
Variation in the GRR repeat sequence and structure. (**A**) Presents a sequence logo built from the alignment of the DGRP repeat units. The height of the letter at each position signifies the frequency of that amino acid in the multiple sequence alignment, and the colour denotes hydrophobicity (blue: hydrophilic, blue: neutral, black: hydrophobic). (**B**) Similarity clustering analysis of GRR2 haplotypes reveals two large groups of *D. melanogaster* haplotypes (gold and red) and one putatively recombinant haplotype (orange). Haplotypes are illustrated using colour-codes for the 16 distinct repeat units corresponding to the arbitrary character identifiers A-P. In some Sanger-sequenced Kenyan haplotypes (labelled ‘Ken’) a repeat unit could not be determined, denoted by a white square. Note that repeat unit L is diagnostic of haplotype group 1, and units D and K are diagnostic of group 2. (**C**) Histogram of the frequency of each haplotype in the DGRP population. Most haplotypes occur at low frequency, with some high and intermediate frequency haplotypes.

Consistent with the antiviral role of *Drosophila* Ago2, the other domains of this protein display strong evidence of positive selection, exhibiting locally reduced diversity around the gene through selective sweeps, and elevated rates of amino acid substitution (Obbard et al, 2006; Obbard et al, 2011; Kolaczkowski et al, 2011). We have previously argued that this rapid adaptive evolution may be driven by virus-mediated selection, through the action of viral suppressors of RNAi (Obbard et al, 2009b), such as those seen in Drosophila C Virus and *Drosophila* Nora Virus (van Rij et al, 2006; van Mierlo et al, 2014). The reportedly high level of variation within the *D. melanogaster* GRR region is therefore surprising, as one might expect diversity to be continually removed by nearby selective sweeps. One possible explanation is that the high diversity and differentiation seen in the GRR is associated with a low constraint on this sequence, combined with high rates of recombination and replication slippage mediated mutations (e.g. Jeffreys et al, 1988). Alternatively, if the GRR domains are involved in the antiviral function of Ago2, or interact with VSRs, the high diversity seen in Ago2 GRRs could reflect the action of diversifying selection—which is a common outcome of many models of host-parasite coevolution. (e.g. Antonovics and Thrall, 1994; Sasaki, 2000)

Whether or not the high divergence and diversity seen in GRR2 is an evolutionary consequence of virus-mediated selection, a virus-related role for GRR2 might be reflected by segregating functional variation associated with GRR2 haplotype. In principle, this could be identified by a genome-wide association study (GWAS) such as that which identified *pastrel* (Magwire et al, 2012). However, as repeat variants are challenging to reconstruct or identify using short sequencing reads (Treangen and Salzberg, 2012), GWAS analyses have largely been limited to SNP and simple structural variation. Thus previous GWAS analyses of viral resistance in *Drosophila* (Magwire et al, 2011; Magwire et al, 2012) have been unable to test for phenotypes associated with highly repetitive sequences, and instead could only have detected its impact through linkage with neighbouring SNPs. But, because SNP diversity is low in *Ago2*-surrounding region, the scale of linkage disequilibrium (LD) is short in *Drosophila*, and LD between a SNP and neighbouring hypermutable loci break down rapidly (Sawaya et al, 2015), a role for GRR variation in determining viral resistance remains untested.

Here we characterise the sequence diversity of the Ago2 GRR region in insects, and use Pacific - Biosciences SMRT long-read sequencing of RT-PCR amplicons to generate full GRR haplotypes for 127 lines of the *Drosophila* Genetic Reference Panel (DGRP; Mackay et al, 2012). We use these data to re-examine the evolution of this domain and its potential role in antiviral defence. In doing so we not only demonstrate the value of long-read technology for performing genome-wide association studies (GWAS) when complex repetitive loci are present, but also illustrate the potential challenges associated with such analysis using short-read technology alone. We provide the first robust *Ago2* GRR haplotypes for natural populations, and identify likely haplotypes in publicly available short read data, and quantify differences in the frequency and composition of GRR haplotypes between African and North American populations. Using published GWAS data (Magwire et al, 2012) to test for an association between GRR haplotype and virus survival phenotypes, we detect a small but nominally significant association of GRR haplotype with longevity of DCV-infected flies. However, we were unable to confirm this association with a second independent experiment using recombinant inbred lines.

## Methods

### Comparison of the GRR across insects

We obtained the GRR repeat unit for other insect species by using tBLASTx with default parameters to query all arthropod RefSeq RNA sequences using the *Ago2* region just C-terminal to the GRR from *D. melanogaster*. We manually selected repetitive sequences as input for Tandem Repeat Finder (v4.07b, Benson, 1999) with a mismatch and indel penalty of 5 and minimum alignment score of 50. The insect reference tree was inferred using MrBayes (v2.13, Huelsenbeck and Ronquist, 2001) with an HKY85 substitution model and gamma-distributed rate variation with invariable sites, using conserved sequences from the original tBLASTx search aligned in MUSCLE (v3.8.31, Edgar, 2004) as input. The high divergence between GRR sequences, including extensive indel variation, makes it extremely challenging to infer positional homology (i.e. alignment) in the GRR regions. We therefore used the frequency feature profile phylogeny building tool (v.3.19, Sims et al, 2009) to quantify similarity between the GRR of insects, as this approach can be used in the absence of alignment. Frequency feature profiles break the nucleotide or amino acid sequence into a distribution of kmers and compares these distributions against each other taking into account similarity between amino acid residues. The frequency feature profiles were constructed in two ways: in the first, GRR repeat unit consensus sequences were used as input to cluster GRRs, and in the second the entire GRR region was used. In each case, the topology of these clusters were compared to the MrBayes tree, using a kmer size which maximised similarity of the feature frequency profile tree to the MrBayes tree, as it is expected that the GRR shares the same history as the rest of Ago2.

### Sample preparation

We sequenced the GRR region from a subset of the *Drosophila* Genetic Reference Panel (DGRP) and 7 other closely related *Drosophila* species. The DGRP constitute a collection of highly inbred lines from *D. melanogaster* collected in Raleigh, NC in 2003 (Mackay et al, 2012) that have previously been sequenced using the Illumina platform to provide a public resource for GWAS. However, as short-read sequencing cannot easily be used to reconstruct repetitive sequences, such as the GRR region of *Ago2*, we generated new amplicon data for the *Ago2* GRR region from 127 of these lines. To avoid sequencing the long intron between GRR1 and GRR2, (RT-)PCR was performed on RNA extracted from 10 flies per line to obtain an amplicon containing the full *Ago2* GRR1 and GRR2 regions. For species other than *D. melanogaster*, sample origins are as described in Longdon et al (2011). For all species, RNA was extracted using Trizol (Ambion) according to the manufacturer’s instructions. Three forward primers were designed separately for the *Drosophila melanogaster/simulans/mauritiana* clade, the *Drosophila yakuba/erecta/santomea* clade, and for *D. ananassae* based on published genome sequences (PCR primer sequences: 15F *D. yakuba*: ATGGGAAAGAAGAACAAATTCAAGG; 30F *D. melanogaster*: GAACAAGAAAGGAGGACAGG; 18F *D. ananassae*: ATATAAGGATGACGGGAAGC). PCRs shared a single reverse primer designed to amplify all species (1550R CAGCTTATCCACCGAGTAGCA) except for *D. ananassae* (GTCGACATTAAGAAACGGTT). Paired barcode sequences from the Pacific Biosciences SMRT Portal v1.4 were added to the 5’ end of each primer, along with the padding sequence GGTAG. Barcoded amplicons were then combined into 10 pools of 16 samples and gel purified for sequencing.

### Long read amplicon sequencing and analyses

Samples were pooled in groups of 16 and subject to Pacific Biosciences SMRT-cell sequencing (NERC Biomolecular Analysis Facility, Liverpool). *D. melanogaster* raw reads were demultiplexed and filtered in the SMRT portal by 5 minimum passes around the circular template, with a minimum predicted accuracy of 70%, a minimum insert size of 1000 bases, and a minimum barcode score of 22. From these, 5-pass circular consensus sequences (5CCS) were called for each read (raw read processing was performed by NERC Biomolecular Analysis Facility, Liverpool). Although these 5CCS reads may still contain errors, to obtain the final consensus sequence for each sample we grouped all 5CCS reads by length, and then removed reads whose length was observed in less than 10 reads. This filtering resulted in a single peak of read lengths for each amplicon (e.g. Figure S1) in all but one fly line. In this one case (DGRP-306), we detected two high-frequency haplotypes, suggesting that this line is heterozygous at the GRR region, and this sample was excluded from all subsequent analyses. Consensus sequences from 5CCS reads within the length class resulted in high-confidence haplotypes from 127 of the DGRP lines, which were used in further analyses. In addition, eight haplotypes from a Kenyan (Nairobi) population, which were previously obtained by Sanger sequencing of long PCR products (Obbard et al 2006), were also included in the analysis. GRR sequences were also obtained from single lines of *D. simulans, D. sechellia, D. mauritiana, D. yakuba, D. santomea, D. erecta*, and *D. ananassae*. These were analysed in the SMRTportal with the parameters described above, but with a minimum insert size of 500 bp. We used BLAST (2.2.31+, Camacho et al, 2008) to assign each long 5CCS read to species based on coding sequence to the 3’ of GRR2, then we grouped reads by species and read length. Peaks in read length were again assumed to be indicative of a distinct amplicon, and analyses were performed as in the *D. melanogaster* samples. To cluster haplotypes (Figure 2) by repeat unit, the distinct repeat units observed in *D. melanogaster* were each labelled with an identifying letter, such that a haplotype can be denoted a string of repeat-unit identifier letters. We then used text-based feature frequency profiles (hash length of 2) to cluster and visualise haplotypes by repeat unit similarity (Sims et al, 2009).

**Figure S1:**
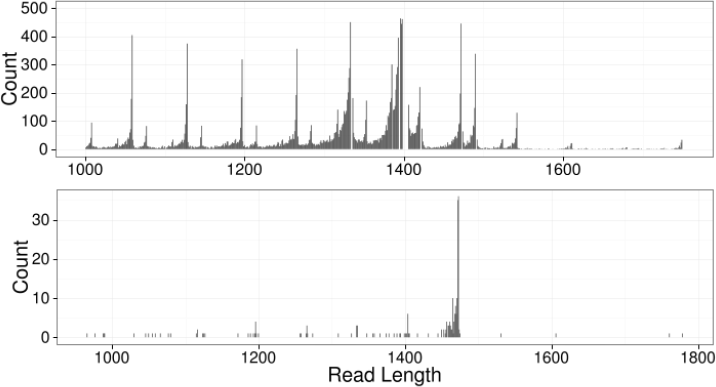
PacBio sequencing reads of the GRR. (*Above*) Read length distribution for all PacBio reads across GRR samples. Multiple peaks confirm that the population is variable in length for this region. (*Below*) An example line sequenced. Consensus sequences were created from reads which had a length supported by more than 10 reads.

### Characterisation of GRR repeats in published short-read data

To explore the utility of published short-read sequencing in the reconstruction of the *Ago2* GRR, we obtained short read sequences of DGRP (Accession number: PRJNA36679, Mackay et al, 2012) and *Drosophila* ‘Nexus’ lines (Lack et al, 2015: Table S1 Accession numbers). To retain reads deriving from the region of interest, all reads were mapped to the full set of 127 sequenced GRR haplotypes using Bowtie2 (v. 2.2.4, Langmead and Salzberg, 2012) with default parameters, retaining all read pairs for which at least one mate mapped. An attempt was made to assemble these reads de novo using Velvet (v1.2.10, Zerbino and Erbino, 2008), using the hash length for each individual that maximised contig length, and using the expected coverage and insert length data from the short read archive.

To assess whether the distribution of repeat units in short read sequences could be used to infer GRR2 haplotypes, we used Jellyfish (v.2.2.3, Marcais and Kingsford, 2011) with a kmer size of 69 (the size of a GRR repeat in *D. melanogaster)* and a lower coverage bound of 2 (although this parameter had no qualitative effects when varied from 0 to 10) to infer counts for known repeat units in each sample. To ensure we only included samples with sufficient coverage of the GRR to reliably infer haplotypes, we filtered out those without reads supporting repeat units GRR2-G and repeat unit GRR2-A, and without ten reads supporting GRR2-E (all of which occurred in all samples, with GRR2-E being most common). The retained samples were then normalised by total read count to obtain a proxy for relative abundance of repeat units in each sample.

### Linkage Disequilibrium analysis

We combined our GRR haplotype data with known SNPs and indels within 5 KB on either side of *Ago2* from the DGRP dataset (http://dgrp2.gnets.ncsu.edu/data/website/dgrp2.tgeno), replacing any reported sequence within the GRR with our own long-read sequence data. We then calculated a multiallelic extension of r^2^ (Hill and Robertson, 1968), which provides an accurate metric of linkage disequilibrium (LD) among multiallelic loci (Zhou et al, 2007). The analysis was performed using our data coded as entire haplotypes (and therefore highly multiallelic), and also as a series of SNPs and indels from alignment of the haplotypes.

The rapid increase in frequency of a beneficial allele is expected to lead to extended regions of high LD around the swept allele (termed ‘haplotype homozygosity’); Sabeti et al, 2002) and to quantify this, we used the program nSL (v.0.47 Ferrer-Admetlla et al, 2014), was used to calculate the nSL statistic. This is similar to the more widely-used iHS statistic (Voight et al, 2006), except that distance is measured as the number of segregating sites rather than map distance, making it more robust to recombination rate variation. Moving along the sequence, at each polymorphic site nSL calculates the average number of consecutive polymorphisms associated with either the ancestral or derived allele in question. Either exceptionally large or small values of the nSL statistic are evidence that a variant has rapidly increased in frequency. For *D. melanogaster* we polarised the sites with the *D. simulans* genome by parsimony, aligned by LastZ (v.1.02.00), and standardised the nSL statistic by allele frequency.

### Association with viral phenotypes and infections

To test whether variation in the GRR haplotype is associated with variation in viral resistance, we used data from previous GWAS studies (Magwire et al, 2011; Magwire et al, 2012) of the DGRP lines for resistance against three different viruses. These were *Drosophila C Virus* (DCV, a horizontally transmitted and highly pathogenic Dicistrovirus naturally infecting *D. melanogaster*; Brun and Plus, 1980; Webster et al, 2015); *D. melanogaster Sigma Virus* (DMelSV: a vertically transmitted Rhabdovirus naturally infecting *Dmel*; Brun and Plus, 1980; Longdon et al, 2012; Webster et al, 2015), and *Flock House Virus* (FHV, a horizontally transmitted Alphanodavirus naturally infecting beetles, closely related to Newington virus of *D. immigrans* (Webster et al, 2016). We fitted general linear mixed models using the R package MCMCglmm (v2.22, Hadfield, 2010) with DGRP line and replicate block (block equivalent to date for FHV and DCV) as random effects, and known segregating functional variants (*pastrel* for *DCV*, and *ref(2)p, CHKov,* and *ge1* for DMelSV) and GRR haplotypes as fixed effects.

The final model was:

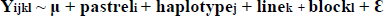

Where μ is the mean survival time and ɛ is a normally-distributed error term. If linkage disequilibrium is sufficiently large, it may be difficult to separate the effect of GRR haplotype from the effect of (partially) linked SNPs. Therefore, to examine whether the GRR haplotype is acting as a marker for a neighbouring causal SNP, we also fitted models in which each flanking SNP was tested for an association with mortality, and then selected those which were nominally significant (with no correction for multiple testing) for inclusion in the model outlined above to verify any observed effect was due to the GRR.

### Recombinant inbred line infections

To further test for an association between *Ago2* GRR haplotype and viral resistance, we experimentally infected recombinant inbred lines from the *Drosophila* Synthetic Population Resource (King et al, 2012) with DCV. We categorised lines by *Ago2* GRR haplotype groups based on presence of short reads containing the repeat units GRR2-L (as a marker for haplotype group 1) or GRR2-D and GRR2-K (as markers for haplotype group 2). The length of the linked region around the GRR region was calculated in each recombinant inbred line, and 100 lines from each haplotype group were selected with the aim of minimising the impact of linked variants (i.e. lines were chosen on the basis of nearby break points). Infections were performed by injecting *Drosophila C Virus* abdominally into 10 flies (per vial, an average of 3 vials per line) at 105 TCID50, chosen on the basis that this dosage caused mortality in approximately one week. Flies were kept at 25 degrees in agar vials and monitored for 7 days post-infection (DPI) with mortality recorded on each day. The data were analysed using a binomial regression in MCMCglmm with the model:

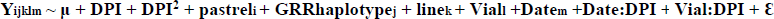

We followed Longdon et al (2011) in coding mortality (Y) as a number of ‘successes’ (the number of flies remaining alive in a vial on a certain day) and ‘failures’ (the number of flies that died on a certain day), as in. This model fits GRR genotype, *pastrel* parent of origin (as a proxy for *pastrel* genotype), and DPI as fixed effects. DPI is encoded as both a linear and quadratic predictor, as mortality tends to decrease after the peak infection. We included DSPR line (genetic background), vial, and date as random effects, allowing for interactions between the DPI and either date or vial effects.

## Results

### Evolution of the GRR across insects

The presence of a GRR region in *Ago2* is conserved across the arthropods, but the GRR evolves extremely rapidly, and the diverse structure of the GRR makes alignment and assembly of these regions challenging. Some species have multiple repeat units, such as *Megachile rotundata* (leafcutter bee)—with repeat units QRRSLAPHG and LKQQQQPLAPQQHHTFA— others have nested repeat units, as in *Tribolium castaneum* (flour beetle), where a region with multiple repeats with consensus QQQWQQQQPQPHP appears to have been duplicated. To circumvent these difficulties, feature frequency profiles (see Materials and Methods) of the GRR and amino acid composition were used to quantify similarity. Conservation of either amino acid composition or repeat unit sequence could imply functional significance of the GRR, and so we examined the GRR of 34 insect species (Figure 1). Feature frequency profiles were constructed from the entire GRR (Figure 1B) or from the consensus repeat unit (Figure S2), and compared to the Ago2 gene tree (Figure 1A, Figure S3). In both cases, the GRR region sequences clustered broadly according to known species relationships but do not reliably reflect more divergent evolutionary relationships. For example, the relationships between *D. melanogaster, D. simulans, D. erecta,* and *D. yakuba* were correctly resolved, but the Drosophilidae did not cluster together in any distance measure. This divergence is in part due to structural differences between GRRs (Figure 1C), as the number and size of repeat units is variable, even between closely related species. Alternatively, amino acid sequence composition is similar across the species analysed, with glutamine the most frequent amino acid residue in all species analysed except *Athalia rosae* (turnip sawfly; Figure 1). This conservation is further illustrated throughout the Drosophilidae (and closest outgroup *M. domestica*), whose GRR is strikingly glycine-rich. These observations argue that although the GRR sequence and structure evolves quickly, the composition may be under selective constraint, implying functionality.

**Figure S2:**
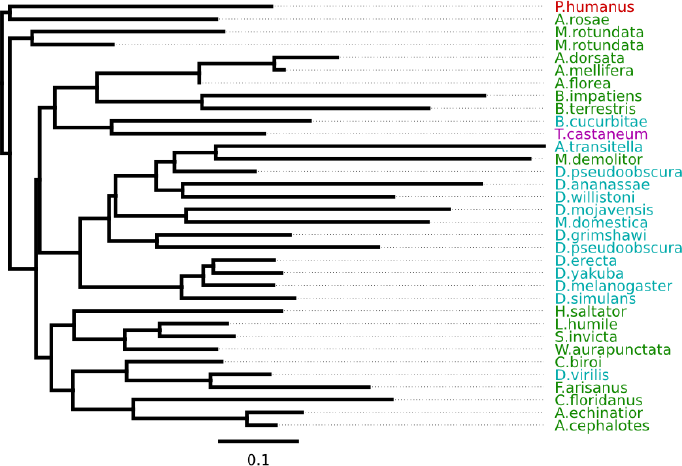
FFP profile clustering from repeat unit consensus sequence. The relationship between consensus sequences for GRR repeat units are unable to recover the true gene tree for *Ago2* (consistent with bottom of Figure 1). Using only repeat consensus sequences should remove the effect of structure (which repeats are next to each other) on this similarity clustering, but reduces the amount of data.

**Figure S3:**
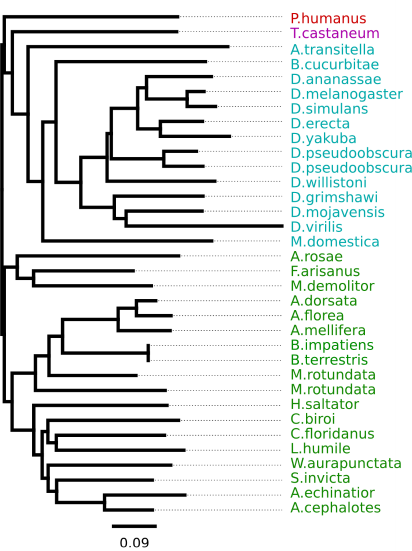
FFP profile clustering from conserved Ago2 sequence. FFP clustering on the conserved Ago2 sequence is mostly in accordance with the maximum clade credibility tree inferred using MrBayes (Figure 1, upper panel). This shows that the FFP clustering approach can infer an approximately correct gene tree from the remainder of Ago2, but not from the GRR region.

### Haplotypes and repeat units in *D. melanogaster Ago2 GRR*

We found extensive repeat polymorphism among the DGRP lines. Among the 127 lines sequenced, we identified three different GRR1 haplotypes and eighteen GRR2 haplotypes. All GRR1 haplotypes comprise one to three perfect repeats of the sequence PQQLQQ, with two repeats being most common (Figure 2). The GRR2 is more complex, with 12 different repeat units (labelled GRR2-A to GRR2-L, Figure 2). The distinct repeat units are all within 3 nucleotide differences of each other and a consensus sequence of GQQQGGHQQGRQGQEGGYQQRPP (Figure 2), and occur 10-15 times in tandem. Most of the GRR2 sequence is composed of two repeat units: GRR2-E (occurring 4-8 times per haplotype) and GRR2-G (occurring 1-6 times per haplotype), which differ at a single amino acid position. In contrast, the majority of repeat units are rare— only occurring in one haplotype, and are most likely the result of recent single base pair mutations (e.g. GRR2-J). Together, the GRR1 and GRR2 alleles form 23 distinct GRR haplotypes in our dataset, which we infer differ from one another by one or two mutation or recombination events (single base changes, whole-repeat insertions and/or deletions, and gene conversion). Clustering GRR haplotypes by repeat unit composition (see materials and methods) identifies two largely distinct haplotypes classes (coloured gold and red in Figure 2), and one putatively recombinant haplotype (*GRR Hap11* – coloured orange) in the DGRP sample. Based on this clustering dendrogram, we have attempted to reconstruct the recent history of the GRR region (Figure 3).

**Figure 3:**
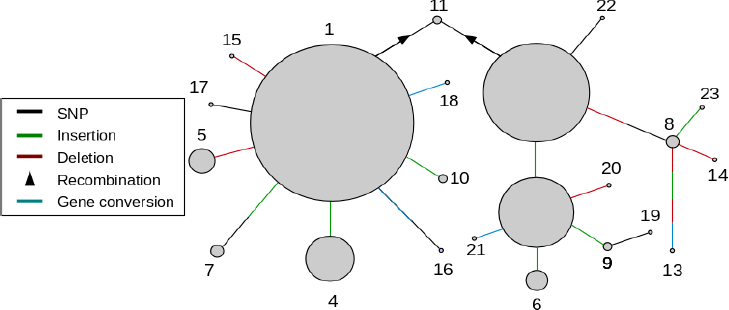
Reconstructed recent history of the GRR. A network showing the inferred relationship between different GRR haplotypes (circles), with circle area corresponding to the frequency in our sample of the DGRP, and connectors representing different mutation or recombination events. Note that there are some haplotypes whose relationship is not easily linked with the others, for example, *GRR Hap13* is unlike any other haplotype sequence and there are large differences between *GRR Hap1* and *GRR Hap2*. In other cases it is not clear whether convergent mutation or recombination produced a particular haplotype - for example, the different GRR1 variants each occur in the background of multiple GRR2 variants (see Figure 2).

Many of these GRR haplotypes occur at a low frequency in this population, with 11 of the 23 haplotypes occurring only once in our sample (Figure 2, Figure 3). There are three high frequency haplotypes (*Haps1 – 3*) with the latter two differing by only one repeat unit. Interestingly, there are many differences between the *Hap1* and *Hap2/Hap3* groups, such that no simple single mutational event could convert one to the other. Further, the haplotypes more closely related to *Hap1* occur at low frequencies and are no more than two mutational events from *Hap1* itself, suggesting they may have been formed recently. This observation is at odds with the high frequency of *Hap1*, and may indicate a recent increase in the frequency of the Hap1 group. In further support of this idea, despite the approximately equal frequency of the Hap1 and Hap2/3 classes in the population, nucleotide diversity in Ago2 and a 100 kb surrounding region is much lower in haplotypes from the *GRR Hap1* clade than those in *GRR Hap2/3*, indicating this *GRR Hap1* is younger than expected given it’s frequency (Figure S4). Nevertheless, there does not seem to be any evidence for significant extended haplotype homozygosity in the remainder of the gene (Figure S5).

**Figure S4:**
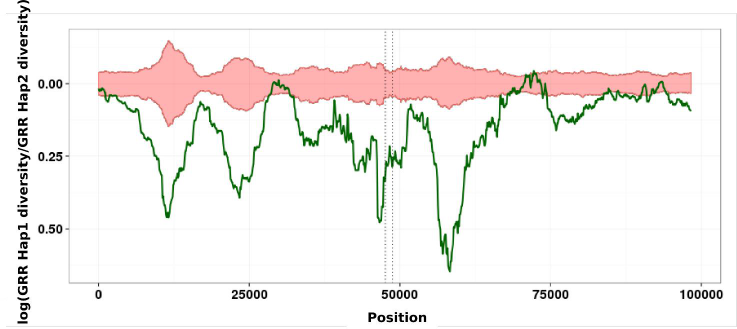
Lower diversity in the GRR Hap1 group relative to GRR Hap2 group. The log of the ratio of *GRR Hap1* clade to *GRR Hap2* clade diversity is plotted (green line) as a sliding window along the chromosome at 50 kb to either side of *Ago2.* The diversity in haplotypes which carry the *GRR Hap1* clade alleles show a surprising lack of diversity in a large area around *Ago2.* The red area shows the extremes of diversity differences expected by chance given the diversity in that region. The dotted lines show the gene region of Ago2.

**Figure S5:**
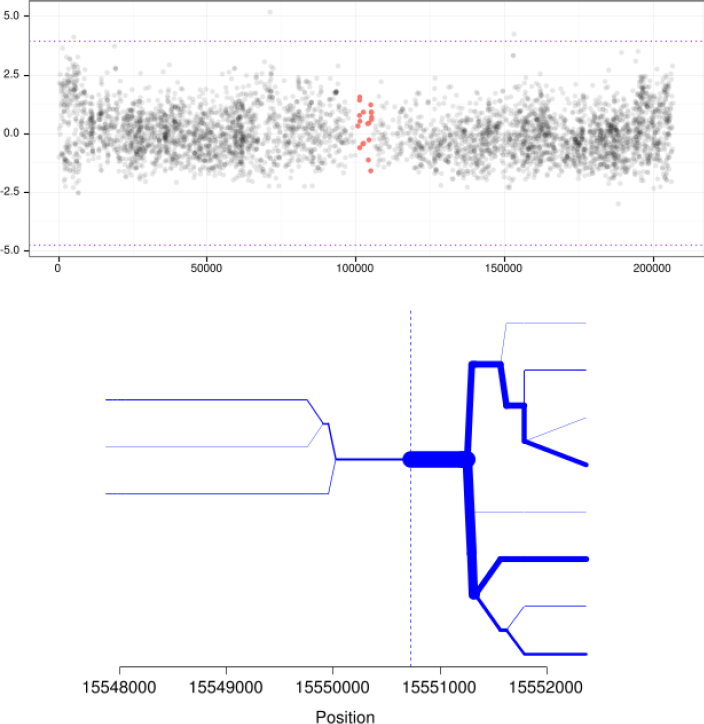
AGO2 nSL signature in the DGRP and haplotype bifurcation diagram. (*Above*) The frequency-standardised nSL statistic at each polymorphic site surrounding *Ago2* indicates no abnormal haplotype structure (red points show polymorphic sites within the *Ago2* gene, black points signify surrounding polymorphic sites). The dotted lines show a threshold at 3 interquartile ranges from the 1st and 3rd quartiles (i.e. extreme outliers). (Below) Visualising the breakdown of haplotype homozygosity supports this conclusion. The bifurcation diagram showing Extended Haplotype Homozygosity was created using the R package rehh (v.1.13, Gautier and Vitalis, 2012). Missing data for the intron upstream of the GRR2 in the DGRP means there is relatively few samples to calculate haplotype homozygosity for this region.

We also analysed 8 Sanger-sequenced GRR2 haplotypes from a Kenyan population of *D. melanogaster* (Obbard et al, 2006) and compared them to the DGRP haplotypes (Figure 2). Notably, 7 of the 8 Kenyan haplotypes were distinct from each other, and in these 7 haplotypes, four new repeat units were found (GRR2-M, GRR2-N, GRR2-O, GRR2-P; Figure 2). This may suggest that the diversity in the DGRP is a subset of African diversity, as expected from the evolutionary history of this species (Lachaise and Silvain, 2004; reviewed in Stephan and Li, 2007). GRR2-L, the defining repeat unit of the GRR Hap1 clade found in the DGRP (gold branches in Figure 2) was rare in this sample of 8 Kenyan sequences, although not absent, suggesting the presence of substantial population structure in GRR.

Although we were unable to reconstruct reliable GRR haplotypes from short-read data, we were able to identify the presence of specific repeat units such as *GRR-L*, which is diagnostic of the *GRR Hap1* clade, and GRR-D andK, which are diagnostic of *GRR Hap2/3* clade. We therefore took advantage of the recent release of the *Drosophila* Genome Nexus, which includes the DGRP as well as individuals sequenced from Africa and France, (Lack et al, 2015) and characterised the distribution of repeat units in these lines (Figure 4). There are repeat units specific to both African (*GRR2-O* and *GRR2-N*) and North American (*GRR2-B, GRR2-H, GRR2-I,* and *GRR2-J*) populations, although those peculiar to North America were all rare variants. French lines also cluster together, characterised by co-occurrence of *GRR2-L* and *GRR2-K* – the defining features of each of the two large classes defined by the DGRP, indicating these lines may be recombinants or heterozygotes. Short read data also suggested that *GRR2-L* is rare in Africa, whereas *GRR-D/K* are common and often co-occur. These observations indicate that the *GRR Hap1* clade has risen in frequency since *D. melanogaster* arrived in N. America.

**Figure 4:**
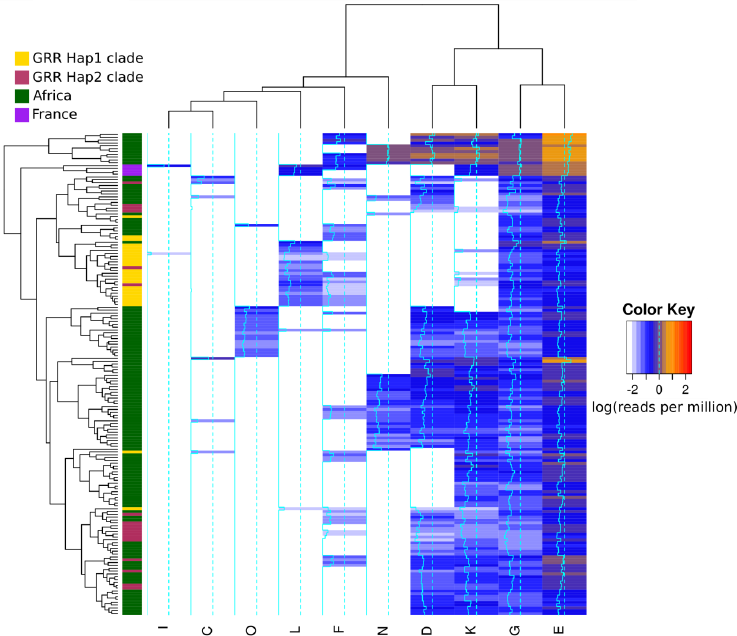
Repeat units in the *Drosophila* Nexus lines. Clustering of the distribution of repeat units in short read data for a sample in *Drosophila* populations taken from the nexus dataset (Lack et al, 2015). Lines were excluded if no short reads were found for ubiquitous repeat units (see materials and methods). *GRR Hap1* clade and *GRR Hap2* clade are those found in the clustering analysis of Figure 2. DGRP *GRR Hap1* clade appears to be derived from an ancestral African population, whereas *GRR Hap2* clade is more divergent, and represents a subset of African diversity. Also, notice the existence of population-specific repeat units (e.g. repeat units *O* and *N*) and population specific co-occurrence of repeats (e.g. repeat units *L* and *K* in France).

### Associations between GRR haplotypes and survival during virus infection

The role of the GRR region is unknown, but as Ago2 is a major effector of antiviral immunity in *Drosophila* (van Rij et al, 2006), it could function during antiviral defence. Using previously published survival data, we found no significant association between GRR haplotype and resistance to Flock House Virus (95% CI for GRR effect: −0.12 – 0.4, *MCMCp* = 0.30) or Sigma Virus (95% CI for GRR effect: −0.15 – 0.03, *MCMCp* = 0.242) infection in the DGRP. However, when fitting GRR haplotype as a fixed effect, we found that *Hap3* alleles increased longevity following challenge with Drosophila C Virus (DCV).by approximately 0.7 days relative to *Hap1* alleles (*MCMCp = 0.012*, [95% CI 0.21 – 1.17] days; Figure S6). This appears to be due to the GRR2 region, as inclusion or exclusion of GRR1 state had no effect. A second model in which GRR1 data were excluded, identified both *Hap2* and *Hap3* as significantly increasing survival relative to Hap1 (*Hap2: MCMCp* = 0.006, 0.56 [ 0.15 – 0.97] days); Hap3: *MCMCp* = 0.006, 0.64 [0.23-1.07] days). However, the observed effect is small relative to the effect of the known resistance variant pastrel^T^ (Magwire et al, 2012), which increases longevity in the same experiment by 2.07 days (MCMCp <0.001, [1.58 – 2.54]).

**Figure S6:**
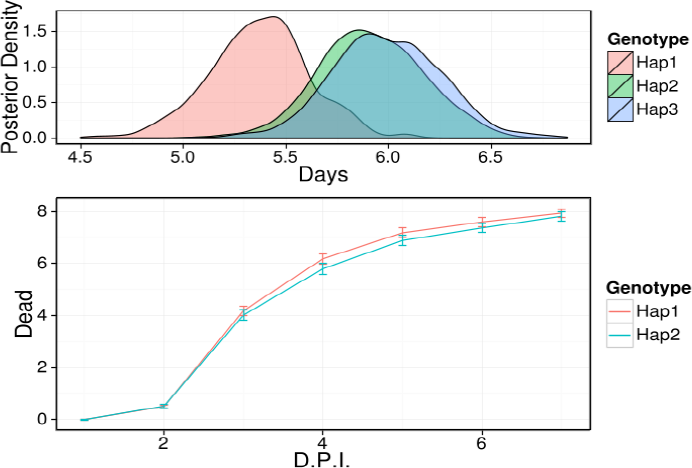
Involvement of GRR during DCV infection. (*Upper*) Estimated posterior density of the survival time (in days) post-infection for *GRR Hap1, GRR Hap2* and *GRR Hap3* using the GWAS data for the DGRP from Magwire et al (2012). *GRR Hap2* and *Hap3* are the two highest frequency haplotypes in the *GRR Hap2* clade (See Figure 2). (*Below*) However, upon infection with DCV, the two haplotype clades show no significant difference in mortality throughout infection in selected DSPR lines.

Given the small size of the effect, multiple tests across viruses, and marginal p-values, we elected to perform a second independent test using a subset of the recombinant inbred lines provided by the Drosophila Synthetic Population Resource (DSPR; King et al, 2012). In this experiment, although mean survival time was again greater for Hap2 than Hap1, this trend was not significant (pMCMC = 0.646; Figure S6). This is unlikely to be due to low power, as we were able to detect a significant association with genotype at the know resistance locus *pastrel* (pMCMC < 0.001). We are therefore unable to replicate the nominally significant effect of GRR haplotype on survival in the DGRP.

## Discussion

### GRR Amino acid composition is conserved, but repeat unit sequence and structure is not

We observe a high degree of sequence divergence in the Ago2 GRR across insect species. Even over very short timescales, there is high divergence in copy number and repeat unit sequence (Figure 1). This could be explained by a high rate of partial inter-repeat replication slippage, causing the creation of new repeat units from the existing ones, and making the sequence unrecognisable in a relatively short period of time (e.g. Dmel and Dsim GRR2 seqeunces, which are unalignable despite only 2.5 My since they shared a common ancestor) In contrast to the sequence of the GRR, we find that the amino acid composition is conserved across the insect species analysed. Based on these observations, we envision a scenario where negative selection acts at the level of amino acids (e.g. to maintain a certain charge or hydrophobicity) but either lack of constraint or positive selection acts at the level of repeat unit sequence and structure.

### GRR repeats are highly variable within *D. melanogaster*, and may be under selection

Repeat number polymorphism in the Ago2 GRR of laboratory lines was previously reported by Hain et al (2010), and our long-read sequencing of a natural population of *D. melanogaster* (the DGRP; Mackay et al, 2012) confirms that this variation is also widespread in the wild. However, our sequencing demonstrates considerable length convergence among haplotypes, such that only 7 different haplotype lengths were present among the 23 distinct haplotypes, and 8 of the 23 distinct haplotypes had the same length (1.035 kbp; Figure 2)—including haplotypes in both the *Hap1* and *Hap2* GRR groups. We found that the haplotypes falling into the *Hap1* clade appear to have recently increased in frequency in the North American (DGRP) population. This is supported by a lower diversity surrounding *GRR Hap1* than *GRR Hap2* clade haplotypes, despite the expectation that neutral diversity in linked regions should scale positively with the frequency of the allele. The increase in frequency of the *GRR Hap1* clade could be due to drift (e.g. during a bottleneck) or selection, such as parasite-mediated selection acting on *Ago2* GRR repeat region itself. However, given the known selective history of *Ago2* (Obbard et al, 2011), this distribution of haplotype frequencies could also be explained by incomplete linkage to a nearby hard sweep carrying GRR Hap1 to a high frequency (e.g. Schrider et al, 2015).

### Ago2 GRR variation is not strongly associated with survival after viral challenge

In other genes, extended low-complexity tracks of a single amino acids have known functions, including been implicated in transcription factor binding (e.g. glutamine, proline, alanine), protein aggregation (glutamine), and cellular localization (histadine), and recently the Q-rich opa repeats of *Notch* have been found to be involved in developmental defects (Gerber et al, 1994; Salichs et al, 2009; Gemayel et al, 2015; Rice et al, 2015). But, although the long-term conservation of the Ago2 GRR among pancrustacea argues that it is maintained by selection, the function of this repeat region remains unclear. As viral suppressors of RNAi (VSRs) have been proposed as the likely drivers of the rapid protein evolution of Ago2 (Obbard et al, 2009), and high diversity is predicted by many models of host-parasite coevolution (e.g. Antonovics and Thrall, 1994; Sasaki, 2000) it is tempting to speculate that the Ago2 GRR may play a role in VSR evasion. For example, the GRR could act to cover residues that underlie Ago2-VSR interactions, or as a bait region, sequestering VSRs away from the catalytic residues of Ago2. However, although *Ago2* GRR showed a significant association with survival after *DCV* infection in our re-analysis of published data from 127 of the DGRP lines, we were unable to replicate this using carefully selected lines from the DSPR. These conflicting results may reflect a false positive from the DGRP analysis, or low power in the DSPR analysis, perhaps due to the challenge inherent in categorising GRR haplotypes using short read data. In either case, it is clear any association must be weak relative to previously identified segregating functional polymorphisms, such as *pastrel*.

### The potential importance of complex repeat sequences in GWAS studies

We find that LD within the GRR, and between the GRR and surrounding variants, is low (Figure S7), indicating that any phenotypic association with this repeat region would be difficult to identify through GWAS using linked sites only. Additionally, the convergence in length between highly divergent GRR haplotypes means that simple length assays may not be suitable to differentiate between haplotypes and may miss important variants. More generally, our study suggests that short read sequencing, such as that currently employed by the majority of association studies, is not a viable option for repetitive regions as we were only able to assemble one correct *Ago2* GRR haplotype among the 117 DGRP datasets using public sequence read data. Clustering by repeat unit presence in short read data confirm our PacBio-sequenced haplotypes (Figure 4), but may only be useful if there is prior knowledge to the possible repeat units in a population and if the region is sequenced in high depth. For example, reads with repeat units *GRR2-*A, *GRR2-G* and *GRR2-E* (which occur in every haplotype) were not always detectable in the short read data for a sample. This indicates that GRR coverage can be low, that incorrect haplotype inference was not only due to assembly errors, and may indicate that the GRR region has unusually low coverage – perhaps because it is not conducive to short read sequencing. Together, these attributes argue that sequencing repetitive regions can provide a depth of understanding not attainable by looking at length variation alone.

**Figure S7:**
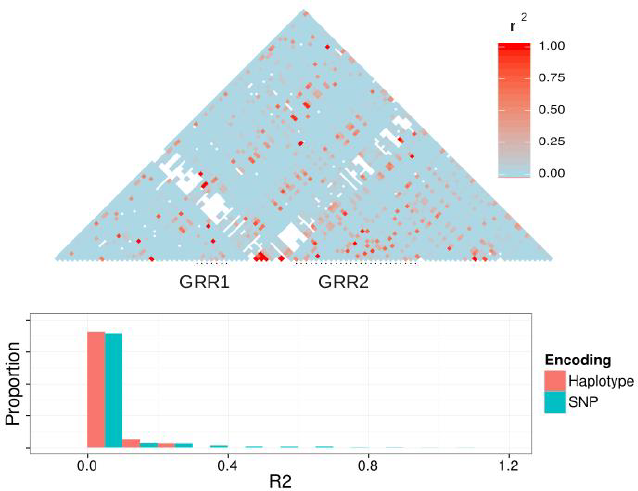
Linkage between GRR and surrounding 10KB and r^2^ values between haplotypes and single GRR SNPs. Pairwise linkage diagram between GRR and surrounding area (above) and a histogram summarising these values. GRR alleles were either broken into a series of SNPs or compared as a single haplotype to calculate r^2^. Linkage is overall low between this region and flanking area, and most information is not recovered by linked SNPs when haplotypes are used to calculate r^2^. As expected, when coding the GRR as one highly multiallelic locus, LD with surrounding areas was much lower, as r^2^ values for low frequency alleles is small and new haplotypes can be continually formed by both recombination and mutation.

## Acknowledgements

We thank Stuart Macdonald for sharing of the DSPR recombinant inbred lines, Jarrod Hadfield for helping with statistical analyses, and Francis Jiggins and Daniel Fabian for making the DGRP virus survival data available to us. PacBio data generation and analysis were carried out by the Centre for Genomic Research which is based at University of Liverpool. This work was funded by a Natural Environmental Research Council Biomolecular Analysis Facility small grant (NBAF 895) to DJO. WHP is funded by the Darwin Trust of Edinburgh, and work in DJO’s lab is supported by a Wellcome Trust strategic award to the Centre for Immunity, Infection and Evolution (WT095831; http://www.wellcome.ac.uk/)

## Data Availability

Haplotype sequences have been submitted to GenBank under the accession numbers KX069093 – KX069218.

## References

1. Antonovics J, Thrall PH. The Cost of Resistance and the Maintenance of Genetic Polymorphism in Host-Pathogen Systems. Proc Biol Sci. 1994;257(1349):105–110. http://www.jstor.org/stable/50298.

2. Benson G. Tandem repeats finder: a program to analyze DNA sequences. Nucleic Acids Res. 1999;27(2):573–580. http://www.pubmedcentral.nih.gov/articlerender.fcgi?artid=148217&tool=pmcentrez&rendertype=abstract. Accessed February 26, 2016.

3. Boland A, Tritschler F, Heimstädt S, Izaurralde E, Weichenrieder O. Crystal structure and ligand binding of the MID domain of a eukaryotic Argonaute protein. EMBO Rep. 2010;11(7):522–527. doi:10.1038/embor.2010.81.

4. Brun P, Plus N. The viruses of Drosophila. In: The Genetics and Biology of Drosophila.; 1980:625–702.

5. Camacho C, Coulouris G, Avagyan V, et al. BLAST+: architecture and applications. BMC Bioinformatics. 2009;10(1):421. doi:10.1186/1471-2105-10-421.

6. Carmell MA, Xuan Z, Zhang MQ, Hannon GJ. The Argonaute family: tentacles that reach into RNAi, developmental control, stem cell maintenance, and tumorigenesis. Genes Dev. 2002;16(21):2733–2742. doi:10.1101/gad.1026102.

7. Edgar RC. MUSCLE: multiple sequence alignment with high accuracy and high throughput. Nucleic Acids Res. 2004;32(5):1792–1797. doi:10.1093/nar/gkh340.

8. Fan H-C, Ho L-I, Chi C-S, et al. Polyglutamine (PolyQ) Diseases: Genetics to Treatments. Cell Transplant. 2014;23(4):441–458. doi:10.3727/096368914X678454.

9. Ferrer-Admetlla A, Liang M, Korneliussen T, Nielsen R. On detecting incomplete soft or hard selective sweeps using haplotype structure. Mol Biol Evol. 2014;31(5):1275–1291. doi:10.1093/molbev/msu077.

10. Garnery L, Vautrin D, Cornuet JM, Solignac M. Phylogenetic relationships in the genus Apis inferred from mitochondrial DNA sequence data. Apidologie. 1991;22(1):87–92. doi:10.1051/apido:19910111.

11. Gautier M, Vitalis R. rehh: an R package to detect footprints of selection in genome-wide SNP data from haplotype structure. Bioinformatics. 2012;28(8):1176–1177. doi:10.1093/bioinformatics/bts115.

12. Gemayel R, Chavali S, Pougach K, et al. Variable Glutamine-Rich Repeats Modulate Transcription Factor Activity. Mol Cell. 2015;59(4):615–627. doi:10.1016/j.molcel.2015.07.003.

13. Gerber H, Seipel K, Georgiev O, et al. Transcriptional activation modulated by homopolymeric glutamine and proline stretches. Science (80-). 1994;263(5148):808–811. doi:10.1126/science.8303297.

14. Hain D, Bettencourt BR, Okamura K, et al. Natural variation of the amino-terminal glutamine-rich domain in Drosophila argonaute2 is not associated with developmental defects. PLoS One. 2010;5(12):e15264. doi:10.1371/journal.pone.0015264.

15. Hill WG, Robertson A. Linkage disequilibrium in finite populations. Theor Appl Genet. 1968;38(6):226–231. doi:10.1007/BF01245622.

16. Huelsenbeck JP, Ronquist F. MRBAYES: Bayesian inference of phylogenetic trees. Bioinforma. 2001;17 (8):754–755. doi:10.1093/bioinformatics/17.8.754.

17. Jorda J, Kajava A V. T-REKS: identification of Tandem REpeats in sequences with a K-meanS based algorithm. Bioinformatics. 2009;25(20):2632–2638. doi:10.1093/bioinformatics/btp482.

18. King EG, Macdonald SJ, Long AD. Properties and power of the Drosophila Synthetic Population Resource for the routine dissection of complex traits. Genetics. 2012;191(3):935–949. doi:10.1534/genetics.112.138537.

19. Kolaczkowski B, Hupalo DN, Kern AD. Recurrent adaptation in RNA interference genes across the drosophila phylogeny. Mol Biol Evol. 2011;28(2):1033–1042. doi:10.1093/molbev/msq284.

20. Kwak PB, Tomari Y. The N domain of Argonaute drives duplex unwinding during RISC assembly. Nat Struct Mol Biol. 2012;19(2):145–151. doi:10.1038/nsmb.2232.

21. Lachaise D, Silvain J-F. How two Afrotropical endemics made two cosmopolitan human commensals: the Drosophila melanogaster-D. simulans palaeogeographic riddle. Genetica. 2004;120(1-3):17–39. http://www.ncbi.nlm.nih.gov/pubmed/15088644. Accessed April 7, 2016.

22. Lack JB, Cardeno CM, Crepeau MW, et al. The Drosophila genome nexus: a population genomic resource of 623 Drosophila melanogaster genomes, including 197 from a single ancestral range population. Genetics. 2015;199(4):1229–1241. doi:10.1534/genetics.115.174664.

23. Langmead B, Salzberg SL. Fast gapped-read alignment with Bowtie 2. Nat Methods. 2012;9(4):357–359. doi:10.1038/nmeth.1923.

24. Lee YS, Nakahara K, Pham JW, et al. Distinct roles for Drosophila Dicer-1 and Dicer-2 in the siRNA/miRNA silencing pathways. Cell. 2004;117(1):69–81. doi:10.1016/S0092-8674(04)00261-2.

25. Lindbo JA, Silva-Rosales L, Proebsting WM, Dougherty WG. Induction of a Highly Specific Antiviral State in Transgenic Plants: Implications for Regulation of Gene Expression and Virus Resistance. Plant Cell. 1993;5(12):1749–1759. doi:10.1105/tpc.5.12.1749.

26. Lingel A, Simon B, Izaurralde E, Sattler M. Structure and nucleic-acid binding of the Drosophila Argonaute 2 PAZ domain. Nature. 2003;426(6965):465–469. doi:10.1038/nature02123.

27. Liu Y, Ye X, Jiang F, et al. C3PO, an endoribonuclease that promotes RNAi by facilitating RISC activation. Science. 2009;325(5941):750–753. doi:10.1126/science.1176325.

28. Longdon B, Hadfield JD, Webster CL, Obbard DJ, Jiggins FM. Host phylogeny determines viral persistence and replication in novel hosts. PLoS Pathog. 2011;7(9). doi:10.1371/journal.ppat.1002260.

29. Longdon B, Wilfert L, Jiggins FM. The sigma viruses of Drosophila. Rhabdoviruses Mol Taxon Evol Genomics, Ecol Cytopathol Control. 2012:117–132.

30. Ma J-B, Ye K, Patel DJ. Structural basis for overhang-specific small interfering RNA recognition by the PAZ domain. Nature. 2004;429(6989):318–322. doi:10.1038/nature02519.

31. Ma J-B, Yuan Y-R, Meister G, Pei Y, Tuschl T, Patel DJ. Structural basis for 5’-end-specific recognition of guide RNA by the A. fulgidus Piwi protein. Nature. 2005;434(7033):666–670. doi:10.1038/nature03514.

32. Mackay TFC, Richards S, Stone EA, et al. The Drosophila melanogaster Genetic Reference Panel. Nature. 2012;482(7384):173–178. doi:10.1038/nature10811.

33. Magwire MM, Bayer F, Webster CL, Cao C, Jiggins FM. Successive increases in the resistance of Drosophila to viral infection through a transposon insertion followed by a duplication. PLoS Genet. 2011;7(10). doi:10.1371/journal.pgen.1002337.

34. Magwire MM, Fabian DK, Schweyen H, et al. Genome-wide association studies reveal a simple genetic basis of resistance to naturally coevolving viruses in Drosophila melanogaster. PLoS Genet. 2012;8(11):e1003057. doi:10.1371/journal.pgen.1003057.

35. Maillard P V, Ciaudo C, Marchais A, et al. Antiviral RNA interference in mammalian cells. Science. 2013;342(6155):235–238. doi:10.1126/science.1241930.

36. Marçais G, Kingsford C. A fast, lock-free approach for efficient parallel counting of occurrences of k-mers. Bioinformatics. 2011;27(6):764–770. doi:10.1093/bioinformatics/btr011.

37. Meister G. Argonaute proteins: functional insights and emerging roles. Nat Rev Genet. 2013;14(7):447–459. doi:10.1038/nrg3462.

38. Obbard DJ, Jiggins FM, Bradshaw NJ, Little TJ. Recent and recurrent selective sweeps of the antiviral RNAi gene argonaute-2 in three species of drosophila. Mol Biol Evol. 2011;28(2):1043–1056. doi:10.1093/molbev/msq280.

39. Obbard DJ, Jiggins FM, Halligan DL, Little TJ. Natural selection drives extremely rapid evolution in antiviral RNAi genes. Curr Biol. 2006;16(6):580–585. doi:10.1016/j.cub.2006.01.065.

40. Obbard DJ, Welch JJ, Kim KW, Jiggins FM. Quantifying adaptive evolution in the Drosophila immune system. PLoS Genet. 2009;5(10). doi:10.1371/journal.pgen.1000698.

41. Obbard DJ, Gordon KHJ, Buck AH, Jiggins FM. The evolution of RNAi as a defence against viruses and transposable elements. Philos Trans R Soc Lond B Biol Sci. 2009;364(1513):99–115. doi:10.1098/rstb.2008.0168.

42. Okamura K, Ishizuka A, Siomi H, Siomi MC. Distinct roles for Argonaute proteins in small RNA-directed RNA cleavage pathways. Genes Dev. 2004;18(14):1655–1666. doi:10.1101/gad.1210204.

43. Parker JS, Roe SM, Barford D. Crystal structure of a PIWI protein suggests mechanisms for siRNA recognition and slicer activity. EMBO J. 2004;23(24):4727–4737. doi:10.1038/sj.emboj.7600488.

44. Ratcliff F, Harrison BD, Baulcombe BC. A Similarity Between Viral Defense and Gene Silencing in Plants. Science (80-). 1997;276(5318):1558–1560. doi:10.1126/science.276.5318.1558.

45. Rice C, Beekman D, Liu L, Erives A. The Nature, Extent, and Consequences of Genetic Variation in the opa Repeats of Notch in Drosophila. G3 (Bethesda). 2015;5(11):2405–2419. doi:10.1534/g3.115.021659.

46. Sabeti PC, Reich DE, Higgins JM, et al. Detecting recent positive selection in the human genome from haplotype structure. Nature. 2002;419(6909):832–837. doi:10.1038/nature01140.

47. Salichs E, Ledda A, Mularoni L, Albà MM, de la Luna S. Genome-wide analysis of histidine repeats reveals their role in the localization of human proteins to the nuclear speckles compartment. PLoS Genet. 2009;5(3):e1000397. doi:10.1371/journal.pgen.1000397.

48. Sasaki A. Host-parasite coevolution in a multilocus gene-for-gene system. Proc Biol Sci. 2000;267(1458):2183–2188. doi:10.1098/rspb.2000.1267.

49. Sawaya S, Jones M, Keller M. Linkage disequilibrium between single nucleotide polymorphisms and hypermutable loci. Genetics. 2016.

50. Schrider DR, Mendes FK, Hahn MW, Kern AD. Soft shoulders ahead: spurious signatures of soft and partial selective sweeps result from linked hard sweeps. Genetics. 2015;200(1):267–284. doi:10.1534/genetics.115.174912.

51. Segers GC, van Wezel R, Zhang X, Hong Y, Nuss DL. Hypovirus papain-like protease p29 suppresses RNA silencing in the natural fungal host and in a heterologous plant system. Eukaryot Cell. 2006;5(6):896–904. doi:10.1128/EC.00373-05.

52. Sims GE, Jun S-R, Wu GA, Kim S-H. Alignment-free genome comparison with feature frequency profiles (FFP) and optimal resolutions. Proc Natl Acad Sci. 2009;106(8):2677–2682. doi:10.1073/pnas.0813249106.

53. Song J-J, Smith SK, Hannon GJ, Joshua-Tor L. Crystal structure of Argonaute and its implications for RISC slicer activity. Science. 2004;305(5689):1434–1437. doi:10.1126/science.1102514.

54. Stephan W, Li H. The recent demographic and adaptive history of Drosophila melanogaster. Heredity (Edinb). 2007;98(2):65–68. doi:10.1038/sj.hdy.6800901.

55. Swarts DC, Makarova K, Wang Y, et al. The evolutionary journey of Argonaute proteins. Nat Struct Mol Biol. 2014;21(9):743–753. doi:10.1038/nsmb.2879.

56. Treangen TJ, Salzberg SL. Repetitive DNA and next-generation sequencing: computational challenges and solutions. Nat Rev Genet. 2012;13(1):36–46. doi:10.1038/nrg3117.

57. van Mierlo JT, Overheul GJ, Obadia B, et al. Novel Drosophila Viruses Encode Host-Specific Suppressors of RNAi. Schneider DS, ed. PLoS Pathog. 2014;10(7):e1004256. doi:10.1371/journal.ppat.1004256.

58. Voight BF, Kudaravalli S, Wen X, Pritchard JK. A map of recent positive selection in the human genome. PLoS Biol. 2006;4(3):e72. doi:10.1371/journal.pbio.0040072.

59. Wang X-H, Aliyari R, Li W-X, et al. RNA interference directs innate immunity against viruses in adult Drosophila. Science. 2006;312(5772):452–454. doi:10.1126/science.1125694.

60. Webster CL, Longdon B, Lewis SH, Obbard D. Twenty Five New Viruses Associated with the Drosophilidae (Diptera). Cold Spring Harbor Labs Journals; 2016. doi:10.1101/041665.

61. Webster CL, Waldron FM, Robertson S, et al. The Discovery, Distribution, and Evolution of Viruses Associated with Drosophila melanogaster. PLoS Biol. 2015;13(7):e1002210. doi:10.1371/journal.pbio.1002210.

62. Wilkins C, Dishongh R, Moore SC, Whitt MA, Chow M, Machaca K. RNA interference is an antiviral defence mechanism in Caenorhabditis elegans. Nature. 2005;436(7053):1044–1047. doi:10.1038/nature03957.

63. Zerbino DR, Birney E. Velvet: algorithms for de novo short read assembly using de Bruijn graphs. Genome Res. 2008;18(5):821–829. doi:10.1101/gr.074492.107.

64. Zhao HH, Fernando RL, Dekkers JCM. Power and precision of alternate methods for linkage disequilibrium mapping of quantitative trait loci. Genetics. 2007;175(4):1975–1986. doi:10.1534/genetics.106.066480.

